# Evidence of target-mediated miRNA degradation in *Drosophila* ovarian cell culture

**DOI:** 10.1101/2023.08.30.555489

**Authors:** Natalia Akulenko, Elena Mikhaleva, Sofya Marfina, Dmitry Kornyakov, Vlad Bobrov, Georgij Arapidi, Victoria Shender, Sergei Ryazansky

**Author notes:** Corresponding author: S.R.

## Abstract

Target-mediated miRNA degradation (TDMD) is a recently discovered process of post-transcriptional regulation of miRNA stability in animals. TDMD is induced by the formation of the non-canonical duplex of Ago-bound miRNAs with the specialized RNA target, and, as suggested for human cell culture, this complex is recognized by the ZSWIM8 receptor protein of the Cullin-RING-ligase complex CRL3. CRL3 ubiquitinates Ago, resulting in proteolysis of Ago and degradation of the released miRNAs. To date, the molecular mechanism of the TDMD process was not studied in other animal species. Here we investigated protein Dora, the *Drosophila* ortholog of ZSWIM8, in the culture of *Drosophila* ovarian somatic cells (OSC). We show that Dora in OSCs localizes in protein granules that are not related to P-and GW-bodies. The knock-out of *Dora* up-regulates multiple miRNAs, including miR-7-5p. Also, we show that Dora associates with proteins of the CRL3 complex, and the depletion of its main component Cul3 up-regulates miR-7-5p. We concluded that the mechanism of TDMD is conserved in humans and *Drosophila*. The knock-out of *Dora* also down-regulates the putative protein-coding targets of miRNAs. One of them is *Tom* from the Brd-C gene family, which is known to repress the Notch signaling pathway. Indeed, in cells lacking Dora, we have observed the down-regulation of *cut*, the marker of the activated Notch pathway. This data indicates that TDMD in OSCs may contribute to modulation of the Notch pathway.

## Introduction

The plethora of mechanisms controlling the expression and activity of miRNAs in animals has recently been expanded by a novel one, called the target-directed miRNA degradation (TDMD) process (Ameres et al. 2010; Shi et al. 2020; Han et al. 2020; Kleaveland et al. 2018; Bitetti et al. 2018; Kingston et al. 2022; Lee et al. 2013; Buck et al. 2010; Libri et al. 2012; Marcinowski et al. 2012; Cazalla et al. 2010; Ghini et al. 2018; Sheng et al. 2023). During TDMD, miRNA is degraded due to its non-canonical binding with the complementary RNA targets within the complex with the Ago protein. In the duplex miRNA pairs with TDMD target along its whole length except for 3-6 mismatched nucleotides in the middle. In humans, hAgo2 with such a duplex gains the specific conformation that is recognized by the receptor ZSWIM8 as part of the Cullin-RING-ligase complex CRL3 (Han et al. 2020; Shi et al. 2020; Sheu-Gruttadauria et al. 2019). CRL3 ubiquitinates Ago, leading to its proteolysis in proteasomes. CRL3 activity is enhanced on neddylation with NEDD8 of its component Cul3 (Merlet et al. 2009). After proteolysis of Ago, the released miRNAs are proposed to degrade in the cytoplasm. However, it is mostly unknown whether the molecular mechanism of TDMD in non-human species is the same.

The key role in the recognition of Ago engaged with the unusual miRNA:TDMD target complex belongs to the ZSWIM8 receptor containing the zinc finger SWIM domain (Han et al. 2020; Shi et al. 2020; Sheu-Gruttadauria et al. 2019). Homologs of ZSWIM8 in *Drosophila* and nematodes, Dora and ebax-1 respectively, also participate in the TDMD mechanism (Shi et al. 2020; Kingston et al. 2022). In S2 cell culture Dora is able to recognize only Ago1-bound miRNAs, but not Ago2-bound small RNAs mostly belonging to endo-siRNAs (Kingston et al. 2022). The bioinformatic search revealed that SWIM8 orthologs are encoded in the genomes of animals, but not in Protozoa, plants, or fungi (Ryazansky and Akulenko 2023). So, the targeted degradation of miRNAs mediated by ZSWIM8 homologs seems to be widely distributed across Metazoa species. The components of the CRL complexes are also very conserved and ubiquitously expressed in animals (Sarikas et al. 2011; Harper and Schulman 2021), which indicates that the mechanism of TDMD may be roughly the same in various species.

Several biological functions of TDMD were identified. In mice, miR-7 destruction during TDMD by binding to long non-coding RNA *Cyrano* accumulates its other target, circular non-coding RNA *Cdr1as* (Kleaveland et al. 2018). TDMD inhibition leads to miR-7-dependent and miR-671-directed slicing of *Cdr1as*, disrupting normal neuronal function. In another case, miR- 29 is degraded when it binds to the lncRNA *libra* in zebrafish and its homolog *Nrep* in mice in certain brain cells (Bitetti et al. 2018). Mutation of miR-29 binding sites in *libra*/*Nrep* led to locomotor defects. The transcripts of many mammalian herpesviruses induce degradation of cellular miR-17, miR-20a, miR-29 and miR-16 by the TDMD mechanism, which is necessary for viral successful infection (Buck et al. 2010; Cazalla et al. 2010; Lee et al. 2013; Libri et al. 2012; Marcinowski et al. 2012). In *Drosophila* embryos, suppression of miRNA-310/313 by TDMD is required for embryonic cuticle development (Kingston et al. 2022). Recent screening of murine miRNAs destabilized by TDMD in different tissues revealed that it can be involved in the regulation of body size and embryonic development (Shi et al. 2023; Jones et al. 2023). The conservative nature of TDMD implies that it might regulate some ancient and general molecular pathways, but it has not been reported yet.

Although TDMD is found in various species (Jones et al. 2023; Shi et al. 2023; Kleaveland et al. 2018; Bitetti et al. 2018; Shi et al. 2020; Kingston and Bartel 2021; Kingston et al. 2022), the molecular details of TDMD are described only for mammals. To highlight the molecular mechanism of TDMD in *Drosophila*, here we investigated the Dora protein, the *Drosophila* ortholog of mammalian ZSWIM8, in OSC cell culture derived from ovarian somatic tissues. We found that in OSCs Dora localizes in cytoplasmic protein granules, associates with components of the CRL3 complex, and represses multiple miRNAs. In addition, we found that knock-out of the *Dora* gene down-regulates many putative targets of miRNAs, including the *Tom* gene. Our data also indicate that TDMD in OSCs might be involved in modulating the Notch signaling pathway.

## Results

### Dora localizes in cytoplasmic protein granules

The Dora protein was previously shown to be involved in the TDMD pathway in embryos and embryonic cell culture S2 of *Drosophila* (Kingston et al. 2022; Shi et al. 2020). To determine whether TDMD is active in other cell types of *Drosophila*, we aimed to characterize Dora in the culture of ovarian somatic cells (OSCs) of *Drosophila.* For this, in OSC expressing *Cas9* (OSC^Cas9+^) we endogenously tagged the *Dora* gene, whose protein product Dora^HA^ fused with the HA-tag at the C-terminus (Figure 1A). The 3’UTR of *Dora^HA^*also contained the blasticidin-resistance gene *bsd* required for the selection of mutant clones in the selective medium. The expression of *Dora^HA^* in blasticidin-resistant cells was confirmed by Western-blotting (Figure 1C). Immunostaining of OSCs with anti-HA antibodies showed that Dora^HA^ was localized in the cytoplasmic granules (Figure 1B). These Dora granules are similar to membraneless protein condensates formed due to the liquid-liquid phase separation process (LLPS) (Hyman et al. 2014). In fact, treatment with the LLPS indicator 1,6-hexanediol disintegrates Dora^HA^ granules (Figure 1D). Ultracentrifugation of the cellular lysate in the sucrose gradient further confirmed that Dora^HA^ is enriched in the high molecular fraction containing ribonucleoprotein complexes (Figure 1E).

**Figure 1.**
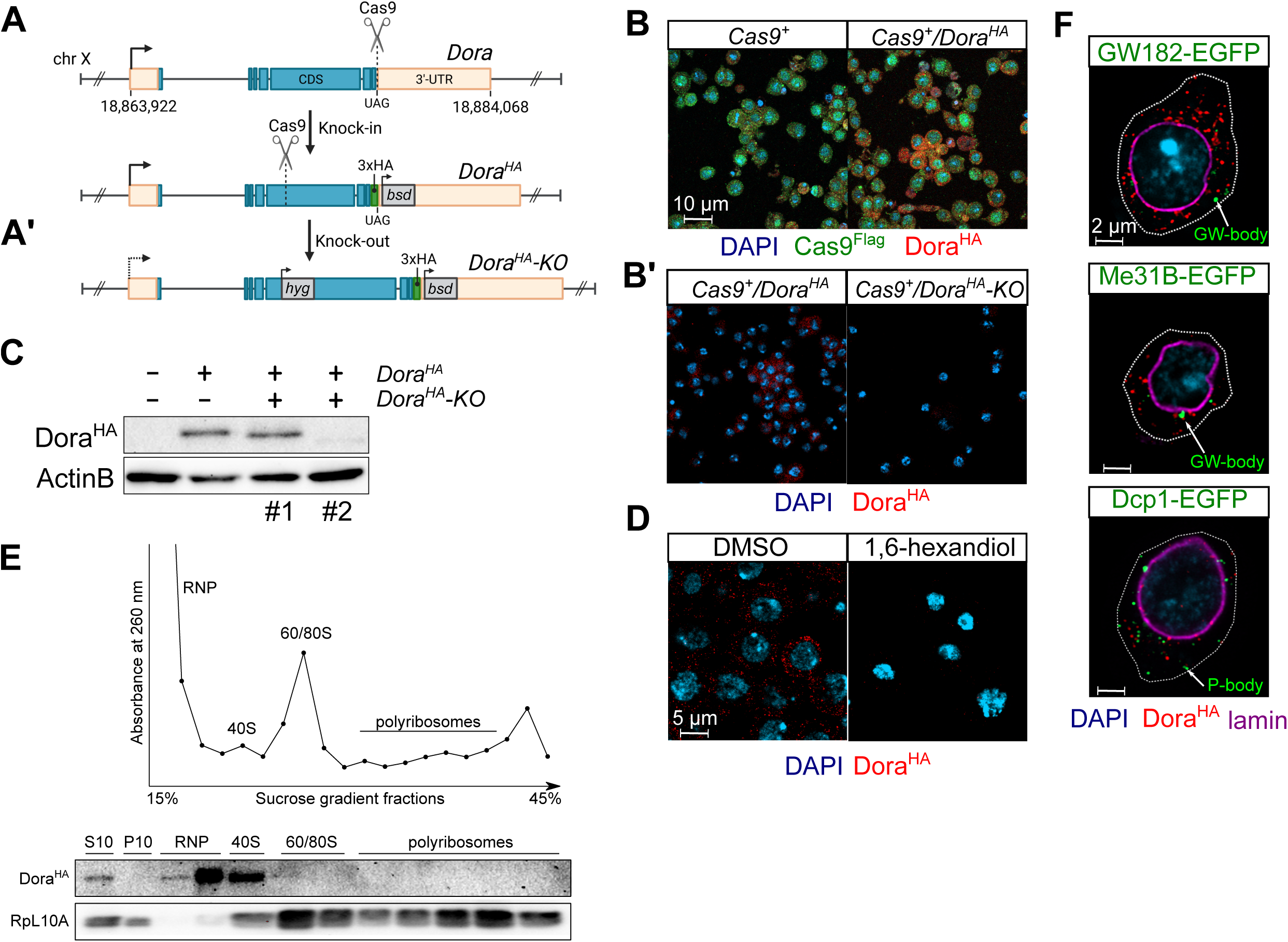
Generation of OSCs with endogenously tagged Dora^HA^ and its subcellular localization. **A.** The scheme of knock-in of endogenous *Dora* using genome editing with Cas9 in OSCs. For knock-in, the 3xHA tag was added at the C-terminus in the frame with the coding region and the blasticidin-resistant gene *bsd* was inserted within 3’UTR. **A’.** For knock-out, the hygromycin-resistant gene *hyg* was inserted into the SWIM domain of *Dora*. **B.** Immunostaining of Dora^HA^ and Cas9^Flag^ in OSC^Cas9+^ and *Dora^HA^* cells with anti-HA and anti-Flag antibodies. **B’**. The immunostaining of Dora^HA^ in *Dora^HA^* and *Dora^HA^-KO* OSCs with anti-HA antibodies. **C.** Western-blot of Dora^HA^ in OSC^Cas9+^, *Dora^HA^*, and *Dora^HA^-KO* cells with anti-HA antibodies. Lanes #1 and #2 are different cellular clones subjected to Cas9-mediated knock-out. Only clone #2 was used for the rest of the work and referred to as *Dora^HA^-KO.* **D.** The anti-HA immunostaining of *Dora^HA^*OSCs treated with 1,6-hexanediol or DMSO as a negative control. **E.** Distribution of Dora^HA^ in the RNP and ribosome fractions. The cellular lysates of *Dora^HA^* OSCs were first centrifuged at 10,000g to obtain the supernatant S10 and the pellet P10 enriched with large membrane organelles. The S10 fraction then underwent ultracentrifugation on a 15% - 45% linear sucrose gradient followed by the fraction collection. The absorbance of fractions at 260 nm is shown on the top panel, while the result of Western-blot with anti-HA or anti-RpL10A antibodies as loading control is shown on the bottom panel. **F.** Immunostaining of Dora^HA^ in *Dora^HA^* OSCs transfected with plasmids encoding *Dcp1-EGFP*, *GW182-EGFP*, or *Me31B-EGFP*.

Then we wondered whether the Dora granules are related to other multiprotein condensates called P-and GW-bodies, which contain many components of the miRNA pathway, including Ago1 (Gibbings et al. 2009; Lee et al. 2009; Miyoshi et al. 2009; Jakymiw et al. 2005; Behm-Ansmant et al. 2006; Eulalio et al. 2007a; Liu et al. 2005b, 2005a; Sen and Blau 2005; Chan and Slack 2006; Standart and Jackson 2007). The typical components of the P-bodies are the Dcp1a and Me31B proteins, while the GW182 protein is enriched in GW-bodies.

To test whether Dora^HA^ granules are related to P-or GW-bodies, we transfected *Dora^HA^* cells with plasmids expressing Dcp1A, Me31B, or GW182 proteins fused with EGFP; the EGFP signal of the fused proteins allows visualization of P-or GW-bodies (Eulalio et al. 2007b). We did not observe any colocalization of Dora^HA^ with the EGFP signal in any case (Figure 1F), indicating that the Dora granules are not related to the P-and GW-bodies.

### Dora represses multiple miRNAs in OSCs

To gain insight into functions of *Dora* in OSCs, we knocked-out *Dora^HA^* by Cas9-mediated insertion of the hygromycin resistance gene *hyg* into its open reading frame (Figure 1A’). The following clonal selection in the selective medium allowed us to obtain a hygromycin-resistant *Dora^HA^-KO* cell line lacking *Dora* expression (Figure 1B’,C). Then we compared the miRNAs repertoires in *Dora^HA^* and *Dora^HA^-KO* cell lines by high-throughput sequencing of the fraction of small RNAs with 21-25 nt in length (Figure 2A). The knocking out of *Dora^HA^* results in differential expression of 17 canonical forms of miRNAs, of which 15 are upregulated (fold change ≥ 2, *p_adj_* ≤ 10e-6) (Figure 2B). The upregulation of several selected miRNAs were also confirmed by the Northern-blot (Figure 2C). The most upregulated miRNAs in *Dora^HA^-KO* compared to *Dora^HA^* cells were miR-10-5p and miR-7-5p (Figure 2A). This is consistent with previous observations that miR-7 is among the main subjects of TDMD in *Drosophila* cell line S2 and several mammalian cell lines (Han et al. 2020; Shi et al. 2020). We also found that for most miRNAs only the guide strand was upregulated in *Dora^HA^-KO* (Figure 2D). This is in agreement with the conclusion that TDMD regulates the abundance of mature miRNAs but not the expression or processing of pre-miRNAs (Han et al. 2020; Shi et al. 2020). The notable exception is miR-10 for which both 5p and 3p strands are upregulated (Figure 2B,D). miR-10 is one of known examples of miRNAs for which both arms of pre-miRNA are almost equally processed (Ruby et al. 2007). This probably suggests that both miR-10-5p and miR-10-3p are regulated by TDMD in OSCs.

**Figure 2.**
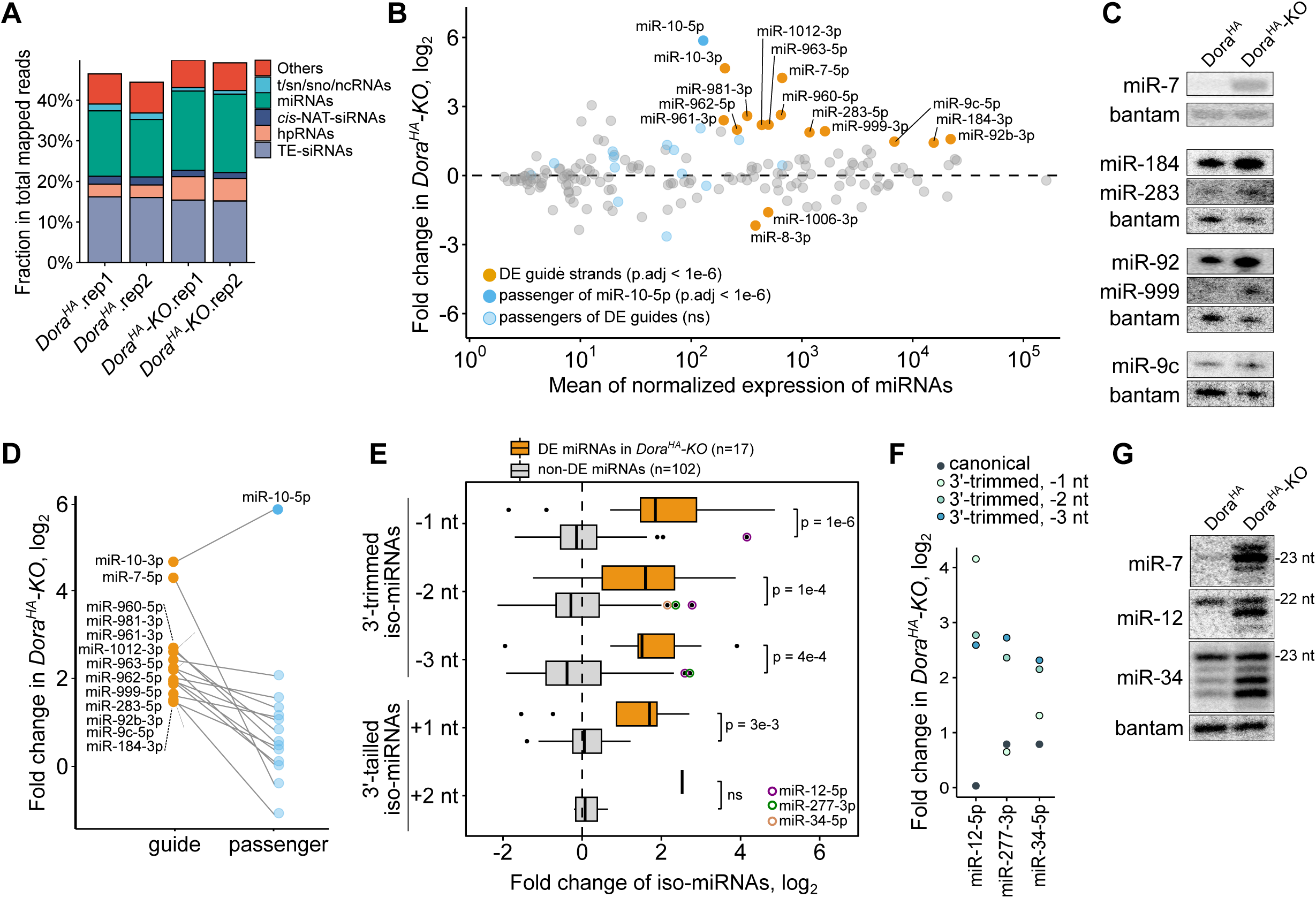
Expression of miRNAs and iso-miRNAs in OSC cells upon knock-out of *Dora^HA^*. **A.** The annotation of 21-25 nt RNAs derived from miRNA, cis-NAT, TE-siRNA, and hpRNA genomic loci. **B.** Differential expression analysis of the canonical forms of miRNAs in *Dora^HA^-KO* relative to *Dora^HA^* OSC cells. For most differentially expressed miRNAs, only their guide strands are affected by loss of *DoraHA* (orange points), while the passenger strands are not (light blue points). For miR-10, both strands are upregulated upon knock-out of *Dora^HA^*, and its minor miR- 10-5p strand is marked by the blue point. **C.** Up-regulation of *Dora^HA^-KO*-sensitive miRNAs was confirmed by the Northern-blot analysis in *Dora^HA^* and *Dora^HA^-KO* cells. The probing to miRNA bantam was used as the loading control. **D.** Comparison of fold change expression (log2-transformed and TMM-normalized values) of 5p-and 3p-strands of the canonical forms of miRNAs that up-regulated upon *Dora^HA^-KO* background. **E.** The barplot shows the fold change expression of iso-miRNAs with one, two or three trimmed (-1, -2, or -3 nt) and tailed by one or two (+1 or +2 nt) nucleotides at the 3’-termini for the upregulated miRNAs (orange bars) and miRNAs that do not change their expression levels (gray bars) upon the knock-out of *Dora^HA^*. The statistical significance was assessed with the Wilcoxon unpaired test. **F.** The fold change expression of 3’-trimmed and canonical forms of iso-miR-12-5p, iso-miR-277-5p, and iso-miR- 34-5p in *Dora^HA^-KO* cells. Only 3’-trimmed iso-miRNAs are upregulated, while their canonical forms are not. **G.** The long-run Northern-blot analysis with probes to miR-7-5p, miR-12-5p, and miR-34-5p demonstrates the accumulation of their trimmed but not canonical forms in *Dora^HA^- KO*. The probing to miRNA bantam was used as the loading control.

The base pairing with the TDMD targets besides the decay of miRNAs also independently triggers the trimming and tailing of their exposed 3’-termini (Han et al. 2020; Shi et al. 2020; Ameres et al. 2010). This TDTT process (after the target-dependent trimming and tailing) becomes prevalent after the impairing TDMD and results in the accumulation of the iso-miRNAs differing from the canonical form of miRNAs by their lengths (Shi et al. 2020). In agreement with this, we observed the accumulation of 3’-trimmed and 3’-tailored iso-miRNAs of *Dora^HA^*-sensitive miRNAs upon knock-out of *Dora^HA^* (Figure 2E). We also noticed that although the abundance of the major forms of miRNA-12, miRNA-277, and miRNA-34 did not change upon *Dora^HA^* knock-out, their trimmed and tailored forms accumulated significantly (*p_adj_* ≤ 10e-6) (Figure 2F,G). Likely, the major forms of these miRNAs are also subjected to TDMD, but in *Dora^HA^* knock-out all of the escaped decay miRNA species undergo TDTT that prevent the accumulation of their mature form.

As previously shown, miRNAs and endogenous siRNAs (en-siRNAs) bound with Ago2 are resistant to the TDMD decay in S2 cell culture (Kingston and Bartel 2021). The fraction of en-siRNAs in embryonic S2 cells is composed mostly of hpRNAs of 22 nt in length processed from long inverted repeat transcripts (Wen et al. 2014). The OCS culture was derived from the somatic cells of *Drosophila* ovaries and in addition to hpRNAs also enriched with Ago2-interacting *cis*-NAT siRNAs and TE-siRNAs with a typical length of 21-22 nt. They are processed from dsRNAs formed due to convergent transcription units that overlap within their 3’-UTRs (*cis*-NAT-siRNA loci) or originated from transposable elements (TE-siRNAs) (Czech et al. 2008; Okamura et al. 2008; Chung et al. 2008; Kawamura et al. 2008). We wondered whether small RNAs predominantly associated with Ago2 are affected in OSCs with knocked-out *Dora^HA^*. Since the genomic location of the *cis*-NAT-siRNA varies in cell lines (Wen et al. 2014), we reannotated them with our own small RNA-seq data. We identified 738 *cis*-NAT loci producing 21-23 nt in length on both genomic strands (Table S2). Then, we remapped and quantified the abundance of 21-23 nt small RNA reads originating from the identified loci, as well as on the hpRNA-generating loci and the canonical sequences of *Drosophila* transposons. We found that *Dora^HA^* loss did not affect the *cis*-NAT-siRNAs or TE-siRNAs (Figure 2A), while the detected numbers of reads corresponded to hpRNAs modestly increased (log_2_ fold change ∼1) (Figure 2A, Figure S1A,B). However, qRT-PCR on one of the most abundant hpRNA esiRNA-sl-1 (encoded in CR46342) did not show any upregulation upon loss of *Dora^HA^* (Figure S1C). Thus, we concluded that similar to S2 cells (Kingston and Bartel 2021), in OSC cells Dora does not regulate the stability of Ago2-bound small RNAs.

### CRL interacts with Dora and is required for the repression of miR-7

During TDMD in humans, the substrate recognition protein ZSWIM8 binds to hAgo2 within the multiprotein ubiquitin-ligase complex belonging to the CRL family (Han et al. 2020; Shi et al. 2020). This complex also contains the scaffold protein Cullin3 (Cul3), the adapter proteins Elongin B (EloB) and Elongin C (EloC), as well as the accessory proteins RBX1 and ARIH1 (Han et al. 2020). This type of CRL complex is unusual since Cullin3 in CRL3 was reported to interact with the BTB adapter protein that also recognizes substrates, while EloB and EloC assist Cul2 in CRL2 and have no their own substrate binding activities (Sarikas et al. 2011). Therefore, we will refer to the CRL involved in TDMD and containing Cul3 and EloB/C as CRL3*. Binding of CRL3* to hAgo2 through ZSIMW8 promotes hAgo2 ubiquitination with subsequent degradation in proteasomes (Han et al. 2020; Shi et al. 2020). The ubiquitin-ligation activity of CRL requires the neddylation of Cullin with NEDD8 (Merlet et al. 2009). To determine whether Dora interacts with CRL in OSCs, we pulled down Dora^HA^ with anti-HA antibodies from the cellular lysate (IP-Dora^HA^) followed by the mass-spectrometry analysis of the immunoprecipitate (Figure 3A,B). For the negative control, we performed mass-spectrometry analysis of proteins bound to anti-HA-beads in the lysate of OSC^Cas9+^ cells without the *Dora^HA^* gene (IP-control). As expected, Dora was the most abundant protein in IP-Dora^HA^ that is almost undetectable in IP-control (Figure 3B). Among other components of the CRL complexes, only EloC was moderately enriched in IP-Dora^HA^, while the EloB and Cullin proteins were not over-represented in the IP-Dora^HA^ (Figure 3B). Applying the criteria of ≤5 ppm of protein abundance signal in the total spectrum of IP-control as well as gaining of ≥15 ppm of excess in IP-Dora^HA^ over IP-control revealed UbcE2M and kl-2 proteins as candidates for interaction with Dora^HA^ (Figure 3B). The UbcE2M protein, also known as Ubc12, catalyzes the attachment of NEDD8 to Cul3, thus enhancing CRL activity (Enchev et al. 2015) and was found as the CRL3* component in humans (Han et al. 2020). Co-immunoprecipitation with EloC and UbcE2M strongly supports Dora as part of the CRL3* complex in *Drosophila*. The lack of overrepresentation of EloB and Cul3 in IP-Dora^HA^ probably is explained by their weak and indirect association with Dora within CRL3*. It is worth noting that we did not reveal any traces of Ago1 in IP-Dora^HA^. We did not observe Ago1 in IP-Dora^HA^ even after pretreatment of OSCs with the proteasomal degradation inhibitor bortezomib, which should prevent proteolysis of Ago1 (Figure S2). Thus, Ago1 dissociates rapidly from CRL3* after ubiquitination.

**Figure 3.**
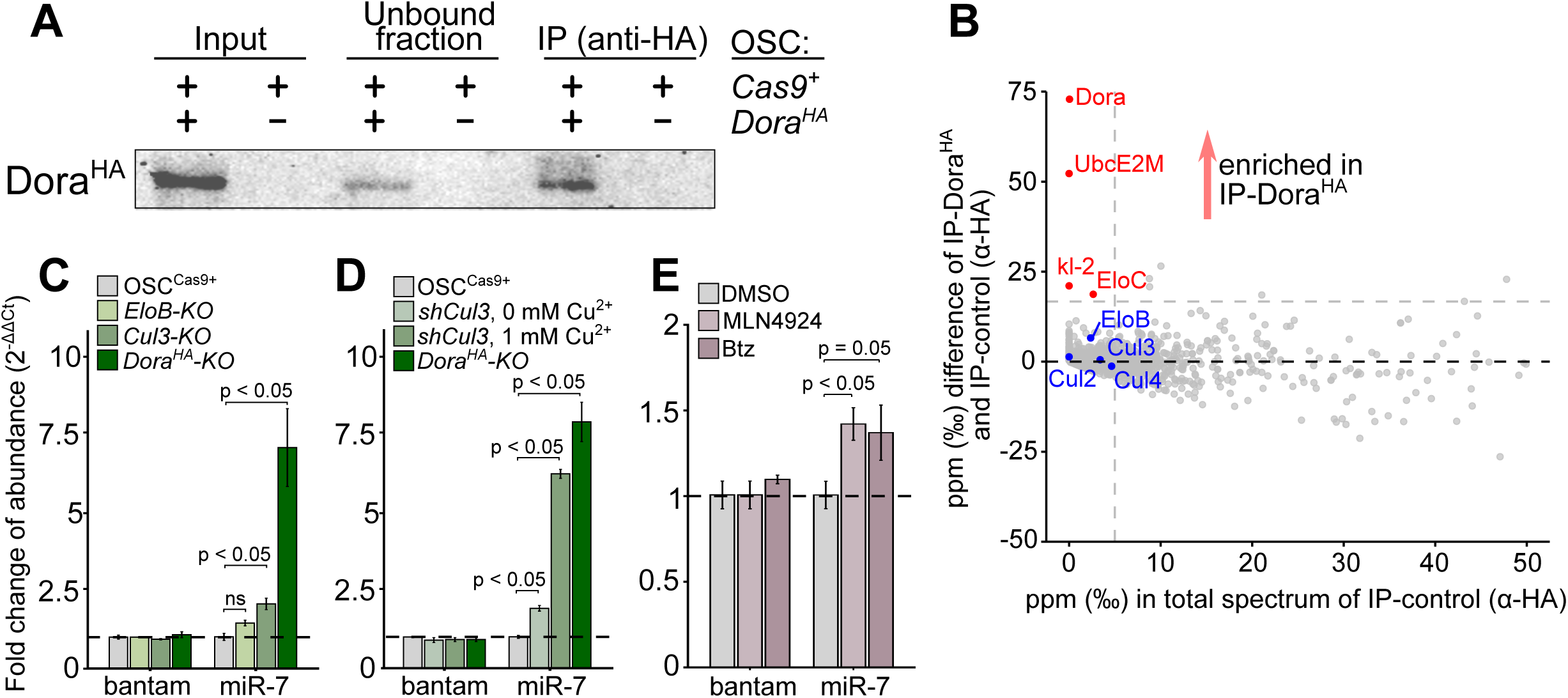
Components of CRL3* interact with Dora and are required for the repression of miR-7. **A.** Immunoprecipitation of Dora^HA^ protein (IP-Dora^HA^) with anti-HA antibodies from the lysate of *Dora^HA^* OSC cells. The anti-HA IP from OSC^Cas9+^ cells was used as the negative control (IP-control). **B.** The result of the mass-spectrometry analysis of the IP-Dora^HA^ in comparison to IP-control. The LFQ intensity for each identified protein was converted to its *per mile* fraction (ppm, ‰) in the total LFQ intensities of all proteins. The result of the subtraction of ppm of IP-Dora^HA^ and IP-control (on the y-axis) was used as a proxy for the enrichment of protein in IP-Dora^HA^. Red points indicate proteins with ≤5 ppm in IP-control but having ≥15 ppm of excess in IP-Dora^HA^ over IP-control. The blue points are the proteins from the CRL family of complexes. **C**. The fold change of abundances of miRNA bantam and miR-7-5p in OSC cells with the knockout of *Cul3*, *EloB,* and *Dora^HA^* genes relative to parental OSC^Cas9+^ cells. The expression levels were measured with qRT-PCR. **D**. The Cu^2+^-induced endogenous knock-down of *Cul3* (*shCul3*, 1 mM of Cu^2+^) accumulates miR-7-5p but not miRNA bantam relative to OSC^Cas9+^. The abundances of miRNAs were measured with qRT-PCR. **E.** The fold change expression of miRNA bantam and miR-7-5p in OSC^Cas9+^ cells (qRT-PCR) treated with MLN4924 or bortezomib relative to control cells treated with DMSO. The error bars in (**C**), (**D**), and (**E**) are the standard errors of the means, and the statistical significance was assessed with the Wilcoxon unpaired test (n=3).

To validate that miRNA repression during TDMD is dependent on CRL3*, we knocked out *Cul3* and *EloB* genes by integrating the hygromycin resistance gene *hyg* into their open reading frames by Cas9 and then measured miR-7-5p expression using qRT-PCR. Genome analysis of puromycin-resistant OSCs revealed that in both cell lines *hyg* was integrated only in one allele of the *Cul3* and *EloB* genes (Figure S3B), and so they are heterozygous in mutations. The failure to obtain homozygous knock-out OSC cultures may be explained by the essentiality of *Cul3* and *EloB* for cell survival. In the *Cul3-KO^+/-^* cells, the expression level of *Cul3* was reduced two-fold (Figure S3A) while miR-7-5p increased modestly but significantly by ∼1.5-fold (Figure 3C). In contrast, in the *EloB-KO^+/-^* cell line, the expression level of *EloB* is reduced by 15 times (Figure S3A) while the expression of miR-7-5p was not changed (Figure 3C). Thus, Cul3 but not EloB may be the component of the CRL3* complex. To further support this finding, we knocked down *Cul3* by shRNA encoding in the genome-integrated cassette. The expression of shRNA^Cul3^ is controlled by the Cu^2+^-inducible metallothionein promoter MT allowing knock-down of *Cul3* on demand (Figure S4A). We have called the system of Cu^2+^-dependent knock-down as the inducible endogenous RNA-interference, and it can be used for efficient repression of any gene that is crucial for cell viability. We have shown that supplementation with 1 mM of Cu^2+^ to cells encoding MT-shCul3 tenfold decreased the expression level of *Cul3* (Figure S4B) while miR-7- 5p accumulates by 6.5 times on a level comparable to that in *Dora^HA^-KO* cells (Figure 3D). This finding is in conjunction with data on knock-out cells that show that Cul3 is required for TDMD in OSCs.

Finally, we tested whether the inhibitors of proteasome activity (bortezomib) and the Cul3 neddylation (MLN4924) affect the level of miR-7-5p. We found that the treatment of OSC cells with bortezomib or MLN4924 leads to the modest by statistically significant upregulation of miR- 7-5p but not bantam (Figure 3E). The association of Dora with CRL3* in OSCs, as well as the requirement of CRL3* and proteasomes for the repression of miR-7-5p, confirms that the molecular mechanism of TDMD in *Drosophila* is similar to that in humans.

### Knock-out of Dora downregulates the targets of miRNAs, including *Tom*

We assumed that the repression of the miRNAs during TDMD enables the expression of their target genes. To identify genes, the expression of which depends on TDMD-mediated miRNA repression in OSCs, we performed RNA-seq capturing 3’-ends of mRNAs in *Dora^HA^* and *Dora^HA^-KO* OSCs (Figure S5). The differential analysis revealed 67 downregulated and 31 upregulated genes in *Dora^HA^-KO* compared to *Dora^HA^* OSCs (log_2_ of fold change ≥ 1.5, *p_adj_* < 10e-3) (Figure 4A). The use of *TargetScanFly* software (Agarwal et al. 2018), predicting the putative miRNAs targets, has shown that 21 (31% of 67) downregulated genes may be targets for miRNAs that are upregulated in *Dora^HA^-KO* (Figure 4B). This proportion is higher than for the upregulated genes, for which only two genes (6% of 31) are the putative targets of the down-regulated miRNAs (*p* < 10e-3, Chi-square test) (Figure 4A, B).

**Figure 4.**
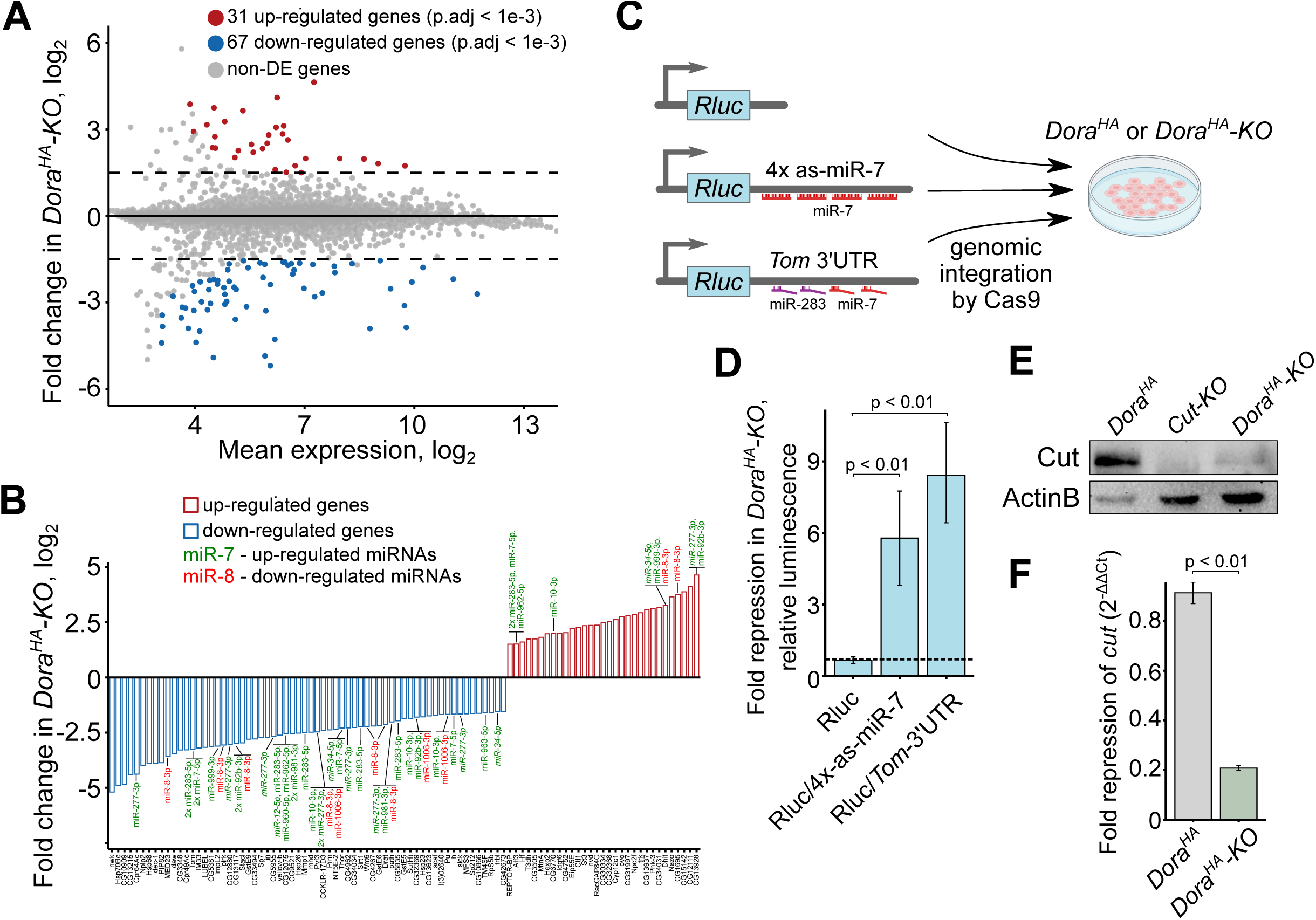
The analysis of gene expression in OSCs with *Dora^HA^* knock-out. **A**. The differential analysis of gene expression revealed by 3’-end mRNA-seq of the OSC transcriptomes in *Dora^HA^-KO* in comparison to *Dora^HA^* cells. **B.** The differentially expressed genes (on the x-axis) are the putative targets of *Dora^HA^-KO* sensitive miRNAs, as well as miR- 12-5p, miR-277-5p, and miR-34a-5p. Target prediction was performed with *TargetScanFly*. **C.** Generation of cell lines with *Dora^HA^* and *Dora^HA^-KO* background that carry the inserted in the genome the reporter constructs encoding *Rluc* with the artificial 3’UTR without any miRNA binding sites (*Rluc*); fused with the 3’UTR of *Tom* (*Rluc/Tom-3’UTR*); or fused with the artificial 3’UTR with four perfectly binding sites for miR-7-5p (*Rluc/4x-as-miR-7*). **D.** The ratio of luminescence of the *Rluc* reporters in *Dora^HA^-KO* relative to *Dora^HA^* cells (repression fold). **E.** The drastic decrease in Cut abundance in *Dora^HA^-KO* and *Cut-KO* cells in comparison to *Dora^HA^* cells was demonstrated with Western blot assay, ActinB was used as a loading control.**F**. The fold repression of *cut* in *Dora^HA^-KO* cells relative to *Dora^HA^* cells and normalized to the expression of *rp49*. The expression level was measured with qRT-PCR. The error bars in (**D**) and (**F**) are the standard errors of the means, and the statistical significance was assessed with the Wilcoxon unpaired test (n=3).

Among the upregulated targets in *Dora^HA^-KO*, there were only two particular genes, *Tom* and *h* (*hairy)*, for which their repression by miR-7 was demonstrated earlier (Stark et al. 2003; Lai 2002, 2005) (Figure 4A). The 3’UTR of *Tom* contains two 8mer binding sites for miR-7-5p and 8mer and 7mer-m8 sites for miR-283-5p, while the 3’UTR of *h* encompasses a single 8mer site for miR-7-5p. To further confirm that *Tom* may be repressed by miR-7-5p and/or miR-283- 5p in OSCs, we engineered the reporter sensor construct, in which the *renilla* luciferase gene *Rluc* is fused with the 3’UTR of *Tom* harboring binding sites for both miR-7-5p and miR-283-5p (Figure 4C). The reporter was inserted at the very same location of the genomes of *Dora^HA^* and *Dora^HA^-KO* OSCs cell lines using Cas9. Measurement of *Rluc*/*Tom-3’UTR* luminescence has shown that it is ∼7x lower in *Dora^HA^-KO* than in *Dora^HA^* reporter-containing cell clones (Figure 4D). The repression level of *Rluc*/*Tom-3’UTR* in *Dora^HA^-KO* was roughly the same as for the other sensor reporter that carried four perfectly matched binding sites for miR-7-5p in its 3’UTR (*Rluc/4x-as-miR-7*). The level of luminescence of the reporter without any miRNA binding sites in 3’UTR was the same in *Dora^HA^* and *Dora^HA^-KO* (Figure 4D). These data are consistent with the increased abundance of miR-7-5p and miR-283-5p in *Dora^HA^-KO* and confirm that *Tom* is the *bona fide* target of one or both of these miRNAs.

Both *Tom* and *h* encode the transcriptional repressors belonging to the Bearded family of genes (Brd-C), and the basic helix–loop–helix (bHLH) family, respectively. These genes are direct targets of the Notch signaling pathway, but Brd-C may also repress Notch through a feedback regulatory loop (Lai et al. 2000). Thus, the miR-7-mediated *Tom* repression in *Dora^HA^- KO* cells may activate the Notch pathway. To further check this possibility, we measured the expression level of *cut* encoding homeodomain protein. The Cut protein is downregulated by the Notch pathway in follicular somatic cells in *Drosophila* ovaries during the middle stages of oogenesis (Sun and Deng 2005). OSC culture was derived from ovarian follicular cells (Niki et al. 2006), so the repression of Cut in OSC may be the indicator of the activated Notch pathway (Stark et al. 2003). We found that in the lack of *Dora^HA^* the abundances of both mRNA and protein products of *cut* are significantly reduced (Figure 4E,G). The abundance of Cut protein in the *Dora^HA^-KO* background dropped to a level similar to control OSCs with knocked-out *cut* (Figure 4G). We concluded that TDMD may repress the Notch pathway by maintaining the high-level expression of its repressor *Tom* by decreasing the stability of miR-7-5p.

## Discussion

Although TDMD is found not only in mammals (Kleaveland et al. 2018; Shi et al. 2020; Han et al. 2020) but also in zebrafish (Bitetti et al. 2018), nematodes (Shi et al. 2020), and *Drosophila* (Shi et al. 2020; Kingston and Bartel 2021; Kingston et al. 2022), the molecular details of TDMD in the latter species are unknown. To highlight the molecular mechanism of TDMD in *Drosophila*, we investigated the Dora protein, the ortholog of mammalian ZSWIM8, in OSC cell culture derived from the ovarian somatic tissues. For this, we generated the OSC line expressing endogenously tagged Dora fused with the HA-tag. We have shown that Dora^HA^ is localized in OSCs in cytoplasmic granules lacking the typical components of P-and GW-bodies (Figure 1D,E,F,G). Thus we suggest that Dora localizes in a novel type of granules. Furthermore, we show that Dora^HA^ was co-immunoprecipitated with the components of CRL3*, namely the adapter protein EloC and the Cul3 neddylation factor UbcE2M (Figure 3B). This result agrees with data on the content of the CRL3 complex involved in TDMD in humans (Han et al. 2020; Shi et al. 2020). It would be interesting to determine in future research whether any component of CRL3* colocalizes with Dora^HA^ in the granules and whether these granules are essential for TDMD in general.

The suppression of TDMD stabilizes multiple miRNAs in variable tissues and cell cultures (Shi et al. 2023; Jones et al. 2023; Han et al. 2020; Shi et al. 2020). Similarly, we revealed the repertoire of miRNAs repressed with the involvement of Dora in OSCs, including miR-7-5p (Figure 2B,C). miR-7-5p is up-regulated on the depletion of *Cul3* encoding the component of CRL3*, and also on the inhibition of both the neddylation of Cul3 and proteasome activity (Figure 3C,D,E). This confirms that miR-7-5p is repressed, at least partly, proteasome-dependently with the participation of CRL3*. Thus, we conclude that our data in *Drosophila* support the mechanism of TDMD in humans.

Among *Dora^HA^-KO*-sensitive miRNAs, miR-10 has both the -5p and -3p arms upregulated, while only the major guide strand of other miRNA is regulated. This likely indicates that some TDMD targets exist for both miR-10 strands. It was shown that the destabilization of one of the miRNA strands by TDMD may facilitate its antisense counterpart to be dominantly loaded and preserved in the complex with Ago and thus may be the mechanism of regulating ‘arm switching’ (Shi et al. 2023; Jones et al. 2023). The control of stability of mature miRNAs and, related to this, the determination of active strands by degradation of their counterparts allow TDMD to affect the content of cellular miRNA pool. The observation that TDMD destabilizes both strands of miR-10 points to the novel dimension of its regulatory potential.

The TDMD targets, in addition to launching the decay of miRNAs, also independently trigger the TDTT pathway during which the exposed 3’-termini of miRNAs are subjected to trimming and tailing (Han et al. 2020; Shi et al. 2020; Ameres et al. 2010). In humans, when TDMD is suppressed, TDTT becomes prevalent, resulting in the accumulation of iso-miRNAs with different sizes (Shi et al. 2020). We have similarly observed the accumulation of iso-miRNAs in *Dora^HA^-KO* OSCs (Figure 2E,F), supporting the interconnections of TDMD and TDTT. Interestingly, for miR-12, miR-277, and miR-34 we have observed only the accumulation of their trimmed forms, but not the canonical ones. There are two alternative explanations. The first one is that there are TDMD targets that differentiate iso-miRs from the major form of miRNAs, and thus TDMD specifically regulates the abundance of iso-miR-12, iso-miR-277, and iso-miR-34 but not their corresponding canonical forms. However, the most plausible explanation is that pairing TDMD targets with the major miRNA forms of miR-12, miR-277, and miR-34 leads mostly to the TDTT pathway instead of miRNA decay. In support of this, miR-34 was reported to be extensively trimmed by 3’-5’ exonuclease Nibbler (Han et al. 2011; Liu et al. 2011). The differentiation between these alternatives requires future research.

Analysis of the transcriptome of *Dora^HA^-KO* cells by RNA-seq revealed that the lack of Dora^HA^ not only stabilized multiple miRNAs but also up-and down-regulated their protein-coding targets (Figure 4A,B). Ago-bound miRNA base pairs to a TDMD target with its seed and 3′ region retaining several mismatched nucleotides in the middle of the duplex. Such duplexes are thermodynamically more stable than the canonical ones provided by only 6-8 nt of the seed. Moreover, the central bulge of the unpaired nucleotides prevents the Ago-cleavage and release of the target strand. This means that the bound TDMD duplex should stick in Ago, locking it down and thus impeding the overall kinetics of the miRNA pathway. One may suggest that the possible functions of the Dora-mediated ubiquitination and proteolysis of Ago1 is the elimination of such locked and nonfunctional Ago1 engaged with TDMD target from the cellular pool. This would make TDMD complementary to the mechanism of the control of quantity and quality of the unloaded Ago (Ryazansky and Akulenko 2023). If so, the knock-out of *Dora* should reduce the fraction of the functionally active Ago1 and derepress miRNA targets. However, we did not observe the massive upregulation of miRNA targets in *Dora^HA^-KO*: only 5 among 31 up-regulated genes are predicted to be targets for *Dora^HA^-KO*-sensitive miRNAs. Thus, the impact of TDMD on the stability of the cellular pool of Ago1 is minimal, and is limited to only a fraction of Ago1 bound with TDMD targets.

Among downregulated genes in *Dora^HA^-KO*, we identified two known targets of miR-7-5p, *Tom* and *h* (Stark et al. 2003; Lai 2002, 2005). The 3’UTR of *Tom*, in addition to two miR-7-5p binding sites, contains two binding sites of miR-283-5p, which is also the subject of Dora-dependent destabilization. *Tom*`s repression by miR-7-5p was demonstrated in the imaginal discs of larvae (Stark et al. 2003; Lai 2002, 2005), and here we also verify this in OSCs (Figure 4C,D). Furthermore, we have observed the down-regulation of the Cut protein that was used as the indicator of activation of the Notch pathway (Figure 4E,G). Both *Tom* and *h* encode the transcriptional repressors belonging to the Bearded family of genes (Brd-C), and the basic helix–loop–helix (bHLH) repressor family, respectively (Lai et al. 2000). The genes from the hHLH and Brd-C families, as well as genes from the E(spl)-C (*enhancer of split*) family, are directly activated by the Notch pathway and one of their functions as transcriptional repressors is to inhibit Notch through a feedback regulatory loop (Lai et al. 2000). Although we do not have direct evidence that Cut is repressed in *Dora^HA^-KO* due to the lack of *Tom*-mediated repression of the Notch pathway, we suggest that this possibility is quite reasonable.

The Notch signaling pathway has a wide range of biological roles in *Drosophila* development. For example, in oogenesis it is required for the differentiation switches of somatic follicular cells in stages 6-7 and 10 (Sun and Deng 2005; Sun et al. 2008). In the differentiation switch on the 6-7 stages, a normal mitotic division is modified to three rounds of endoreplication with a skipped M cell cycle phase resulting in 16 copies of genomic DNA. The proper entry of the follicle cells into the endocycle is initiated by the Notch-dependent downregulation of Cut (Sun and Deng 2005). The endoreplication is also dependent on the miRNA activity since *dicer1* and *pasha* mutations inactivate the Notch pathway, possibly because of the upregulation of miRNA target Delta (Poulton et al. 2011). We proposed above that in OSCs the expression of *cut* depends on *Tom*, which negatively regulates Notch and is targeted by TDMD-destabilized miR-7-5p. If this is correct, then the intriguing possibility is that the entry into the endocycle of follicular cells in ovaries may also be initiated by the down-regulation of TDMD following the stabilization of particular miRNAs (e.g., miR-7-5p) allowing the Notch pathway to repress Cut. It is worth mentioning, that although in ovaries the differentiation shift results in a skipped M phase, we did not find evidence of the inefficient M phase in *Dora^HA^-KO* cells, as the abundance of Cyclin A protein was not affected (Figure S6). Therefore, the entry to the endocycle in ovaries should be promoted by some other factors besides Cut down-regulation. miR-7-5p was shown to repress many Notch-regulated genes from the E(spl)-C and Brd-C families (Lai 2005). It will be interesting to determine whether the activity of *Tom* or other genes of Brd-C and E(spl)-C are permitted by TDMD in follicular cells or other cell types. In general, this paves the way for future research on Notch pathway regulation and the biological functions of TDMD in *Drosophila*.

## Materials and Methods

### Data availability

Raw data of the high-throughput sequencing of miRNAs and mRNAs are available as NCBI BioProject (PRJNA1007640).

### Cell culture

The culture of *Drosophila* ovarian somatic cells OSC (Niki et al. 2006) (kindly provided by Dr. M. Siomi, University of Tokyo) was grown at 25°C in Shields and Sang M3 insect medium (Sigma-Aldrich, #S3652) supplemented with 10% heat-inactivated fetal bovine serum (FBS) (Gibco, #10270106), 10% fly extract, 10 µg/ml insulin (Sigma-Aldrich, #I9278), 0.6 mg/ml L-glutathione (Sigma-Aldrich, #G6013), 50 units/ml penicillin and 50 g/ml streptomycin. To prepare the fly extract, 1 g of 3-7 day-old flies were homogenized in 6.8 ml of M3 medium and centrifuged for 15 min at 1,500 g. The supernatant was heated for 10 min at 60°C and cleaned with centrifugation at 1,500 g for 1.5 h at 4°С. The obtained extract was filter-sterilized through 0.22 µm filters before usage.

### OSCs with stably expressing Cas9

The OSC cells were transfected with the pAc-sgRNA-Cas9^Flag^ plasmid (Addgene plasmid #49330; a gift from Ji-Long Liu) (Bassett et al. 2014); the transfection mix containing 30 µl of serum-free medium M3, 0.5 µg of plasmid, and 2 µl FuGENE-HD (amounts are given per well of a 24-well plate), was incubated for 15 min at RT and added to cells. The selection of OSC^Cas9+^ cells with plasmid integrated into the genome was performed on the medium with puromycin (Gibco, #A1113803) in the concentration of 1 µg/ml.

### Bortezomib, MLN4924 and 1,6-hexanediol treatment

Cells of 2-3 days growth with approximately 70-90% confluency have been treated with Bortezomib (Merck, #504314) at 100 nM for 42 h, MLN4924 (Cell Signaling, #85923) at 100 nM for 70 h. As a control, the treatment of DMSO at 0.1% v/v was used. The optimal concentrations and time of treatment by bortezomib and MLN4924 was chosen in preliminary experiments. For 1,6-hexanediol treatment cells were treated with 2 µg/ml digitonin with or without 10% 1,6- hexanediol (Sigma, #240117) in Shields and Sang M3 medium at 25°C for 1 h.

### Generation of endogenously HA-tagged *Dora* in OSCs

#### The construction of sgRNA producing plasmid and the preparation of donor dsDNA

To obtain endogenously HA-tagged *Dora* in OSC cells, we have applied the Cas9-based approach suggested by (Kunzelmann et al. 2016; Böttcher et al. 2014), in which the donor dsDNA fragment for the homologous recombination is the PCR product containing shoulders complementary to regions near the site of the integration. The sgRNA sequence required for the addition of HA-tag sequence at the C-terminus of Dora was selected with *TargetFinder* (Gratz et al. 2014); the corresponding oligonucleotides (Table S1) were annealed and cloned in the pU6-BbsI-chiRNA vector linearized with *BbsI* (Addgene plasmid #45946; a gift from Melissa Harrison & Kate O’Connor-Giles & Jill Wildonger) (Gratz et al. 2013); the resulting sgRNA^Dora^ plasmid was used as the source of guide RNA. The donor dsDNA was generated by PCR using a pMH4_HA plasmid as a template and primers containing on their 5’-ends 60 nt arm sequences complementary to upstream and downstream regions of the sgRNA cleavage site (Table S1). The pMH4_HA plasmid was constructed on the basis of the pMH4 vector (Addgene plasmid #52529; a gift from Klaus Förstemann) (Kunzelmann et al. 2016), in which Flag-tag was replaced with a 3xHA-tag sequence. The obtained donor DNA fragment contained the sequence of HA-tag and the blasticidin resistance gene *bsd*.

#### Cas9-mediated mutagenesis of OSCs

The *Dora* gene was endogenously HA-tagged in the genome of OSC constitutively expressing *Cas9* (OSC^Cas9+^). First, we inhibited the NHJR process by the knock-down of *lig4* and *mus308* genes. For this, the corresponding dsRNAs prepared according to (Kunzelmann et al. 2016) were added to final concentration of 1 μg/ml of each dsRNA in a serum-free M3 medium to cells having 50-60% level of confluency. After 5 h the medium was supplemented with nutrients and cells were then transfected with the same double-stranded RNAs using FuGENE-HD Transfection Reagent (Promega, #E2311): the transfection mix containing 30 µl of serum-free medium M3, 0.5 µg of each dsRNA, and 3 µl FuGENE-HD (amounts are given per well of a 24-well plate), was incubated for 15 min at RT and added to cells. Three to four days after transfection, half of the cells were transferred to a fresh medium. On the next day cells having 50-60% confluency were transfected with the mix containing 20 µl serum free M3 medium, 375 ng of the plasmid expressing sgRNA^Dora^, 75 ng of the dsDNA donor, and 2 µl FuGene HD (for one well of a 24-well plate). Three to four days after the transfection, the growth medium was replaced with the selective medium with 10 µg/ml of blasticidin S (Applichem, #A3784). The presence of the HA-tag at the C-terminus of Dora in blasticidin-resistant cellular clones was determined by PCR with one primer to HA-tag and another one to the *Dora* gene (Table S1) and by Western-blot analysis with anti-HA antibodies (see below).

### Knocking out of genes

The open reading frame of the *Dora^HA^* gene (SWIM domain) was disrupted with the Cas9-mediated insertion of hygromycin B resistance gene *hyg*. The technique was the same as for the generation of endogenously HA-tagged *Dora*. As a template for the PCR synthesis of the donor fragment, we used pMH4 plasmid in which *bsd* was replaced with *hyg*. The replacement was performed by Gibson assembly (NEBuilder HiFi kit, NEB) of two PCR fragments containing *hyg* from pcDNA5.0-TO plasmid and pMH4 vector without *bsd* (all primer sequences in Table S1). The site for sgRNA-mediated cleavage was chosen by *TargetFinder* and the corresponding oligonucleotides are listed in Table S1. The selection of hygromycin-resistant clones was performed on the medium with 200 µg/ml of hygromycin B (Calbiochem, #400050). The knock-out of *Cul3*, *EloB*, and *cut* was performed in the same way in OSC^Cas9+^; the oligonucleotides designed to synthesize the corresponding sgRNA plasmids are listed in Table S1.

### Western-blotting

About 2×10^7^ of OSCs were lysed in 10 mM Tris-HCl (pH 6.8), 100 mM NaCl, and 7 M urea. About 30 µg of total cell protein was separated on 8% SDS-PAGE and blotted onto Immobilon-P PVDF membrane (Millipore). The membrane was blocked in 0.2% I-Block (Tropix), incubated with primary antibodies for 1 h at RT, and washed three times for 5 min with a PBT buffer (PBS with 0.1% Tween-20). The used primary antibodies are mouse anti-HA (1:1,000, 6E2, Cell Signaling, #2367), rabbit anti-RpL10 (1:1,000, Sigma), mouse anti-Cut (1:200, DSHB, 2B10), mouse anti-ActinB (1:1,000, Abcam, AB8224), or goat anti-CyclinA (1:100, Santa Cruz, N-15). Then, the membrane was incubated for 1 h at RT with secondary AP-conjugated anti-mouse or anti-rabbit antibodies (1:20,000, Sigma), washed three times for 5 min with PBT and once for 15 min in 20 mM Tris-HCl (pH 9.8) and 150 mM NaCl. The chemiluminescence was developed with Immun-Star AP Substrate (Bio-Rad) with subsequent visualization using the ChemiDoc MP imaging system (Bio-Rad).

### Immunostaining of OSCs

Immunostaining was performed according to the previously described protocol (Ilyin et al. 2017). Cells attached to a coverslip were fixed in 4% formaldehyde in PBS for 12 min and rinsed with PBT (PBS with 0.1% Tween-20) three times. After the permeabilization with PBTX (PBS with 0.1% Tween-20 and 0.3% Triton X-100) for 20 min, cells were blocked with PBTX containing 3% normal goat serum (Invitrogen) for 1 h at RT and incubated with primary antibody in PBTX with 3% normal goat serum for 3 h at RT. After washing in PBTX three times for 10 min, cells were incubated with secondary Alexa Fluor antibodies (1:500, ThermoFisher Scientific) in PBTX with 3% normal goat serum for 1 h at RT and then washed in PBTX three times for 10 min in a dark chamber. The coverslips were then mounted with a drop of SlowFade Gold Antifade reagent (Invitrogen) containing DAPI. The confocal microscopy was performed using an LSM 510 META system (Zeiss) and LSM 900 Confocal with the Airyscan2 super-resolution detector (Zeiss). The following primary antibodies were used for immunostaining: rabbit anti-FLAG (1:100, D6W5B, Cell Signaling, #14793), mouse anti-FLAG (1:150, Sigma, #3165), mouse anti-HA (1:300, 6E2, Cell Signaling, #2367), rabbit anti-HA (1:500, C29F4, Cell Signaling, #3724).

### Co-localization of Dora granules with P-and GW-bodies

OSC cells were transfected with plasmids pAc5.1B-EGFP-DmMe31B, pAc5.1B-EGFP-DmDCP1 or pAc5.1B-EGFP-DmGW182 (Addgene plasmids #21682, #21684 and #22419, respectively; a gift from Elisa Izaurralde) (Eulalio et al. 2007b) as described above and in 48 h cells were immunostained with anti-HA antibodies.

### Sucrose gradient of Dora^HA^

About 2×10^7^ cells were treated with cycloheximide CHX (0.1 mg/ml) for 10 min at 25°C, and washed with cold PBS with the same CHX concentration. Washed cells were lysed in 0.5 ml of buffer A (20 mM HEPES (pH 7.6), 100 mM KCl, 5 mM MgCl_2_, 2 mM DTT, 1% NP-40, 0.1 mg/ml heparin, RNase and protease inhibitors, 1 mg/ml CHX). To form a linear gradient, 5 ml of 15% sucrose solution was carefully layered on 5 ml of 45% solution (w/w, in buffer B: 20 mM Hepes (pH 7.6), 100 mM KCl, 10 mM MgCl_2_, 1 mM DTT, 1% NP-40, 1 mg/ml heparin, RNase and protease inhibitors, 0.1 mg/ml CHX). The sealed centrifuge tube was laid horizontally at RT for 2 h, returned to the vertical position, and kept at 4°C for at least 30 min. The lysate was centrifuged at 10,000 g for 15 min; the obtained supernatant and pellet were designated as S10 and P10, respectively. S10 extract was layered on the gradient solution with subsequent ultracentrifugation at 36,000 rpm for 2 h at 4°C in a P40ST bucket rotor (Himac). Fractions of 0.5 ml were manually collected (20-21 in total), precipitated with 10% of trichloroacetic acid, and dissolved in 150 µl (1st fraction) or 50 µl (remaining fractions) of 7 M urea solution; 3 µl of the fractions 1-4 and 10 µl of the fractions 5-20 were mixed with SDS gel loading buffer with subsequent Western-blotting analysis of samples with anti-HA or anti-RpL10 antibodies.

### Northern-blotting of small RNAs

Total RNA from 2×10^7^ cells was purified with RNazol-RT (MRC) according to the manufacturer’s instructions. 10-30 µg of the fraction enriched with small RNAs was separated on denaturing 12% PAGE (8M urea, 0.5x TBE) with 0.5x TBE as electrophoretic buffer (pre-electrophoresis at 10 mA for 20 min), soaked in 0.5x TBE for 15 min and transferred onto the Amersham Hybond-N+ membrane (Cytiva) at 400 mA for 40 min. RNA was immobilized on the membrane with UV crosslinking at 120 mJ/cm^2^ and additional baking for 30 min at 80°C. The membrane was pre-hybridized in HS buffer (100 mM Na-P buffer (pH 7.5), 0.5 M NaCl, 0.1% Ficoll-400, 0.1% Polyvinyl pyrrolidone, 150 µg/ml herring sperm DNA (Promega), 50 µg/ml heparin, 25 mM EDTA, 1% SDS) in the presence of 25% formamide for 1 h at the hybridization temperature. The probes (Table S1) were ^32^P-labeled with T4 PNK (Promega) and purified using Oligo Clean and Concentrator columns (OCC-5, ZymoResearch). LNA or unmodified oligonucleotide probes were hybridized overnight at 50°C or 37°C, respectively, in the HS buffer but without herring sperm DNA. After hybridization, a membrane was washed three times with 2x SSC and 0.2% SDS at the hybridization temperature for 15 min and exposed overnight to a storage phosphor screen with subsequent scanning by Typhoon FLA 9500 (GE Healthcare).

### Small RNA sequencing and bioinformatic analysis

To deplete 2S ribosomal RNA, 10 µg of small RNA fraction extracted with RNAzol RT (MRC) was separated in 15% denaturing PAGE and a gel piece containing 20-25 nt in length RNA was excised. RNA was eluted from the gel overnight in 400 µl of 0.4 M NaCl with rotation at RT and precipitated with 96% ethanol in the presence of linear acrylamide (NEB).

Approximately 100 ng of purified RNA was used for library preparation according to the protocol with randomized splint ligation (Maguire et al. 2020). The sequences of the used oligonucleotides are in Table S1. After PCR amplification by universal P5 and index P7 primers, the libraries were treated with ExoSAP-IT (Promega) and purified with the DNA Clean and Concentrator kit (DCC-5, Zymo Research) to rid of any traces of primers and free adapters. The libraries were sequenced in two replicas for each sample as SE50 on NovaSeq 6000 at Novogene. As a result, we got 12.5-16.5⋅10^6^ of raw reads in each of the four libraries. After the trimming of the 3’-adapters from the reads, the quantification of miRNA species was performed with *bcbio-nextgen* (Chapman et al. 2021) and *mirtop* software (Desvignes et al. 2020, 2018). Differential expression was analyzed with the *isomiRs* package of R (Pantano and Escaramis 2022).

### Annotation of *cis*-NAT-siRNAs producing loci

The calling of loci that produced *cis*-NAT siRNAs was performed on merged four libraries of small RNAs derived from *Dora^HA^* and *Dora^HA^-KO* cells. The reads in range length of 21 to 23 nt were further mapped on the *dm6* genomic assembly of *D. melanogaster* with *bowtie* v. 1.3.1 requiring no mismatches (-v 0 -m 1 --best --strata). All reads corresponding to rRNA, miRNAs, transposons, and sn/sno/tRNAs regions were removed. For the identification of *cis*-NAT siRNA loci, we used *ShortStack* software v. 3.8.5 (Johnson et al. 2016) (--dicermin 21 --dicermax 23 -- pad 50 --mismatches 1 --mmap u --nohp --mincov 20) that outputted 2506 regions. The lowly-expressed regions with <30 reads were removed. Additionally, as *cis*-NAT siRNA loci generate siRNAs from both strands, we further required that the ratio of abundance of reads derived from sense and antisense strands was >0.2 and <0.8. After extending the regions by 100 bp on both sides, the adjacent loci separated by <250 bp were then merged. The identified 738 cis-NAT siRNAs loci are listed in Table S2. The list of hpRNA loci was derived from (Czech et al. 2008; Okamura et al. 2008; Chung et al. 2008; Kawamura et al. 2008), and the corresponding sequences were fetched from the RNAcentral database. For the identification of 21-23 nt TE-siRNAs we used the canonical transposon sequences (https://github.com/bergmanlab/drosophila-transposons, v. 9.41). Raw reads of 21-23 nt in the length of each library were remapped on the specified loci requiring ≤1 mismatch to estimate the abundance of miRNAs and endo-siRNAs, and requiring ≤3 mismatches for TE-siRNAs. The cpm (counts per million) normalization of abundances was performed on the total amount of 20- 25 nt reads mapped on the genome, excluding reads mapped to rDNA loci.

#### mRNA-seq and bioinformatic analysis

The fraction of long RNA was isolated according to the RNAzol RT protocol (MRC) and treated with DNAse I (Ambion). For the transcriptome quantification, we applied a custom 3’-end mRNA-seq library preparation protocol inspired by the QuantSeq FWD kit (Lexogen) (Corley et al. 2019). For this, 150 ng of RNA was mixed with 40 ng of SID-X oligonucleotide (where X = 1, 2, 3 or 4), denatured at 85°C for 3 min followed by cooling to 35°C and kept at this temperature for 2 min. The SID-X oligonucleotide anneals at the poly-A tail of mRNA and contains 8 nt UMI part and 4 nt sample barcode part (Table S1). For RNA from different samples, SID-X (X = 1, 2, 3, or 4) oligonucleotides with sample-specific barcodes were used. After preheating the reverse transcription premix at 35°C, SuperScript II (Invitrogen) was added and the reaction continued at 42°C for 15 min. The short reaction time is required for reverse transcription of only the 3’-end of mRNAs instead of the entire transcript. To prevent SID-X primer hybridization at non-poly-A sites, cooling of the reverse transcription reaction mix lower than room temperature should be avoided. Reverse transcription was terminated with 3 µl of 0.5 M EDTA and then four samples tagged with different SID-X were pooled. The RNA:cDNA duplexes were purified with DCC-5 and eluted with water with subsequent alkaline hydrolysis of RNA with 250 mM NaOH at 65°C for 15 min. Single-stranded cDNAs were purified with OCC-5 and synthesis of the second strand of cDNA was performed. For this, cDNA was mixed with 1 µg of Random_hex_2 oligonucleotide (Table S1), denatured at 98°C and cooled to 25°C; the annealing at this temperature was carried out for 30 min. Synthesis was performed with a Large Fragment of Pol l (NEB) at 30°C for 1 h. To remove excess primers, the reaction mixture was treated with ExoSAP-IT (Promega) at 37°C for 30 min and purified with the DCC-5. Double-stranded cDNA was amplified by PCR with universal P5 and index P7 primers (Table S1) and products of 300- 550 bp in length were cut from the agarose gel and purified with the GeneJET gel extraction kit (Thermo Scientific). The traces of adapter dimers were depleted using AMPure XP beads (Beckman) with a 0.8:1 ratio to gain the final library. This library containing two replicas for each sample was sequenced as PE150 on NovaSeq 6000 at Novogene.

The raw reads were mapped on *Drosophila* transcriptome (Ensemble, v.6.32.107) with *STARsolo*, v. 2.7.10a (Kaminow et al. 2021), while the following demultiplexing of the BAM file for the individual samples according to the sample barcodes and collapsing of the PCR duplicates based on the UMI was performed with *UMI-tools*, v.1.1.2 (Smith et al. 2017) and *bamtools*, v.2.5.1 (Barnett et al. 2011). The metagene profile was constructed with *deeptools*, v.3.5.1 (Ramírez et al. 2016); the annotation was done using the GenomicRange package (v.1.52) and Ensembl GTF (v.6.32.107) in the Biocondutor/R environment (v.4.3.1). The differential expression analysis was done with *DESeq2*, v.1.40.1 (Love et al. 2014) by applying the LFC shrinkage with *apeglm* method (Zhu et al. 2019).

### qRT-PCR measurement of miRNAs and enRNA-sl-1 expression

About 500 ng of small RNA fraction extracted with RNAzol RT (MRC) without ribosome RNA depletion was processed as in the case of small RNA library preparation, but omitting the stage of 5’-adapter ligation. The obtained cDNA was used as a template for qPCR with common primer P7B.7 and miRNA-or enRNA-sl-1- specific primers (Table S1). The Ct values of miR-7, and enRNA-sl-1 were normalized to miRNA bantam (ΔCt). The fold change expression was calculated as 2^-ΔΔCt^ in comparison to their expression in the OSC^Cas9+^ or *Dora^HA^* cells.

### qRT-PCR measurement of gene expression

About 1.5 µg of total RNA extracted from 10^6^ cells with Extract RNA reactive (Evrogen) was subjected to the DNAse I treatment (Ambion) following the reverse transcription with reverse transcriptase Mint (Evrogen) and random primers. The quantitative real-time PCR of cDNA was performed on the DT-96 (DNA-technology) with gene-specific primers (Table S1). The Ct values of *Cul3*, *EloB*, and *cut* were normalized to *rp49* (ΔCt). The fold change expression of genes was calculated as 2^-ΔΔCt^ in comparison to their expression in the OSC^Cas9+^ or *Dora^HA^*cells.

### Mass-spectrometry analysis

#### Immunoprecipitation of Dora^HA^

About 2×10^7^ of OSC^Cas9+^ (IP-control) or *Dora^HA^* (IP-Dora^HA^) cells were resuspended in the culture media without trypsin treatment, collected by centrifugation at 300 g for 5 min, washed three times with cold PBS and lysed in 1 ml of 1x Cell Lysis Buffer (Cell Signaling, #9803) in the presence of a protease inhibitor cocktail (Sigma-Aldrich, GE80-6501-23) for 30 min on ice. The lysed cells were sonicated (amplitude 30%, three repeats of five cycles of 1 sec “ON” / 1 sec “OFF”) with subsequent centrifugation at 16,000 g for 15 min at 4°C. 15 µl of magnetic beads coupled with anti-HA antibodies (Pierce, #88837) were washed three times with PBS and incubated with the cell extract for 1 h at RT under the rotation. After incubation, the beads were washed three times with 200 µl of PBST, and bounded proteins were eluted with 100 µl of the Sample buffer (10 mM Tris-HCl (pH 8.0), 8 M Urea, 2 M Thiourea) at RT for 10 min with manual mixing. The aliquots of input, flow through, and eluate were analyzed by Western blotting with anti-HA antibodies. Protein concentrations in eluates were then measured for each sample using a Bradford Kit and then, eluates were subjected to mass-spectrometry analysis.

#### In-solution trypsin digestion

In eluates from immunoprecipitations in solubilization buffer (8 M Urea, 2 M ThioUrea, 10 mM TRIS-HCl (pH=8)), disulfide bonds of proteins in each sample were reduced with DTT (final concentration 5 mM) for 30 min at RT. Afterwards, Iodoacetamide was added to a final concentration of 10 mM. The samples were incubated in RT for 20 min in the dark, with the reaction being stopped by the addition of 5 mM DTT. Samples were then diluted with 50 mM Ammonium Bicarbonate solution to reduce urea concentration to 2 M. Trypsin (Promega; Cat#V5111) was added at the ratio of 1:100 w/w and the samples were incubated for 14 h at 37°C. After 14 h the reaction was stopped by the addition of Formic acid up to a final concentration of 5%. Finally, the tryptic peptides were desalted using SDB-RPS membrane (Sigma-Aldrich), vacuum-dried, and stored at -80°C before LC-MS/MS analysis. Before LC-MS/MS analysis samples were redissolved in 5% Acetonitrile (ACN) with 0.1% Trifluoroacetic acid (TFA) solution and sonicated.

#### LC-MS/MS analysis

Mass-spectrometry analysis of all samples was performed using the Q Exactive HF mass-spectrometer. Samples were loaded onto 50-cm columns packed in-house with C18 3 μM Acclaim PepMap 100 (Thermo Fisher Scientific), using Ultimate 3000 Nano LC System (Thermo Fisher Scientific) coupled to a mass-spectrometer (Q Exactive HF, Thermo Fisher Scientific). Peptides were loaded onto the column thermostatically controlled at 40°C in buffer A (0.2% Formic acid) and eluted with a linear (120 min) gradient of 4-55% buffer B (0.1% formic acid, 80% ACN) in buffer A at a flow rate of 350 nl/min. Mass-spectrometry data were stored during the automatic switching between MS1 scans and up to 15 MS/MS scans (topN method). The target value for MS1 scanning was set to 3×10^6^ in the range of 300-1200 m/z with a maximum ion injection time of 60 ms and resolution of 60,000. Precursor ions were isolated at a window width of 1.4 m/z and fixed the first mass of 100.0 m/z. Precursor ions were fragmented by high-energy dissociation in a C-trap with a normalized collision energy of 28 eV. MS/MS scans were saved with a resolution of 15,000 at 400 m/z and at a value of 1×10^5^ for target ions in the range of 200–2000 m/z with a maximum ion injection time of 30 ms.

#### Protein identification

Raw LC-MS/MS data from the Q Exactive HF mass-spectrometer were analyzed with MaxQuant (version 1.6.10.43) against protein sequences from FlyBase, a *Drosophila melanogaster* genome database service (version r6.43), with the addition of possible trypsin digestion contaminants. For this procedure, we use the following parameters: Orbitrap instrument type, tryptic digestion with one possible missed cleavage, fixed modification for carbamidomethyl (C), variable modifications for oxidation (M) and acetyl (protein N-term), LFQ label-free quantification. From the list of the identified proteins the contaminants (such as keratins) and highly abundant housekeeping proteins (actin, tubulin, Gapdh, histones, ribosomal proteins, etc.) having >10^7^ LFQ in the samples were excluded (Table S3). Then, LFQ intensities of the remaining 1,145 identified proteins were quantile-normalized between samples and converted to ppm (‰), relative to the total summa of all LFQ intensities for each sample. Only proteins with ≤5 ppm in IP-control but having ≥15 ppm of excess in IP-Dora^HA^ over IP-control were considered as over-represented in IP-Dora^HA^.

### Inducible endogenous knock-down of *Cul3*

To create the inducible knock-down of *Cul3* we constructed the cassette encoding shRNA to *Cul3* under the control of Cu^2+^-inducible metallothionein promoter (MT) and integrated it into the genome with Cas9 (Figure S5). This cassette is based on the pWallium20 vector, where the expression of shRNA is controlled by the minimal *hsp70* promoter and GAL4-activated UAS element (Ni et al. 2011; Perkins et al. 2015). The UAS/*hsp70* promoter in pWallium20 was replaced with MT promoter from pMT-OsTIR1-P2A-H2B-AID-EYFP plasmid (a gift from Christian Lehner), while the *mini-white* gene was replaced with the blasticidin resistance gene *bsd* under the control of the *copia* promoter. The MT-shRNA-bsd cassette was further inserted with Cas9 into the puromycin resistance gene *PuroR* within the *pAc-sgRNA-Cas9^Flag^* integrated into the genome of the OSC^Cas9+^ line. For this, the designed oligonucleotides for sgRNA^puro^ (Table S1) were cloned into the pU6-BbsI-chiRNA vector as described above. To make the donor for the homologous recombination, the cassette MT-shRNA-bsd was cloned by Gibson assembly into *PuroR* of the pAc-sgRNA-Cas9^Flag^ vector with the deletion of most *Cas9* to shrink the plasmid size; the final donor plasmid is designated as pAc_MT_shCul3_bsd. The regions of the *PuroR* gene flanking the cassette MT-shRNA-bsd in the pAc_MT_shCul3_bsd are the shoulders for homologous recombination during the Cas9-mediated genome editing. The detailed scheme of the cloning is available under request and will be published elsewhere.

### *Rluc* reporter constructs

The reporter construct containing the *renilla* luciferase gene (*Rluc*) fused with 3’UTR carrying target sites of miR-7 was integrated into the genomic *PuroR* of OSC^Cas9+^ line. The sequence of 3’UTR was chemically synthesized (four perfectly complementary to miR-7 regions separated with short spacers, 4x-as-miR-7) or PCR amplified from the genome (*Tom*-3’UTR, primers are in Table S1) and cloned into the psiCHECK-2 vector (Promega) linearized with *XhoI* and *NotI* restriction enzymes. psiCHECK-2 with cloned 3’UTR was used as the template for the PCR amplification (primers are in Table S1) of the fragment containing SV40 promoter, *Rluc*- 3’UTR, and polyA site. The donor plasmid was obtained by Gibson assembly of this fragment and plasmid pAc_MT_shCul3_bsd treated with *XmaI* (both to linearize plasmid and remove of MT-shCul3 region from it). We also constructed the control donor plasmid encoding *Rluc* without miR-7 binding sites. The donor plasmids (Rluc/4x-as-miR-7, Rluc/*Tom*-3’UTR and control Rluc) were integrated using Cas9 and sgRNA^puro^ into *PuroR* gene within the *pAc-sgRNA-Cas9^Flag^* genomic cassette of *Dora^HA^*and *Dora^HA^-KO* cell lines with subsequent clone selection on the blasticidin containing selective medium. The luciferase luminescence in the cellular extracts was measured with Dual-Luciferase Reporter Assay (Promega, #E1910) on the Modulus Microplate Reader (Turner BioSystems). The integration of *Rluc* reporters into the same genomic locations (*PuroR* gene) in the different cell lines allows the correct comparison of *Rluc* expression in them.

## Acknowledgments

We thank Christian Lehner for the plasmid pMT-OsTIR1. The research was supported by the Russian Science Foundation (grant No. 22-24-00519 to S.R.). A part of the work related to immunostaining was funded by the Russian Science Foundation grant No. 19-14-00382 to E.M.

## Author contributions

N.A. performed most of the experiments, including the molecular cloning, Western-and Northern-blots, immunoprecipitation, sucrose gradient, qRT-PCR, and library preparation for RNA-seq; E.M. performed all work with cell culture and immunostaining of cells; S.M. performed the molecular cloning, Western-blot, and qRT-PCR; D.K., V.B. performed the molecular cloning; V.S., G.A. performed the immunoprecipitation and mass-spectrometry; S.R. performed the analysis of NGS and mass-spectrometry data; N.A., S.R. conceived and designed the project and edited the draft of the manuscript; S.R. wrote the manuscript with inputs from N.A. and E.M. All authors have read and agreed to the published version of the manuscript.

## Supplementary Information

**Figure S1. The abundance of hpRNAs upon loss of *Dora^HA^*. A.** The distribution of length and normalized abundances (cpm) of sRNA mapped on major sources (CR32205/7, CR18854, CR46342) of hpRNAs in *Dora^HA^* and *Dora^HA^-KO* cells. **B.** The secondary structures of precursors of hpRNAs and the abundance of endo-siRNAs generated from them. The boxes on the structures indicate the positions of the en-siRNAs (bold font) the normalized abundance of which are shown on the barplots. **C.** The result of qRT-PCR on enRNA-sl-1 (CR46342) (Kawamura et al. 2008) and miRNA bantam in *Dora^HA^-KO* cells relative to *Dora^HA^* cells with normalization on miRNA bantam (2^-ddCt^). The error bars are the standard errors of the means.

**Figure S2. The mass-spectrometry analysis of the immunoprecipitate with anti-HA antibodies from the cellular lysate of *Dora^HA^* OSCs treated with bortezomib. A.** The result of the mass-spectrometry analysis of the IP-Dora^HA^ from OSCs treated with bortezomib in comparison to IP-control (IP with anti-HA from *Cas9^+^* cells). The LFQ intensity for each identified protein was converted to its *per mile* fraction (ppm, ‰) in the total LFQ intensities of all proteins. The difference in ppm of IP-Dora^HA^ and IP-control (y-axis) was used as a proxy for the enrichment of protein in IP-Dora^HA^. Red points indicate proteins with ≤5 ppm in IP-control but having ≥15 ppm of excess in IP-Dora^HA^ over IP-control. The blue points are the proteins from the CRL family of complexes. **B.** The comparison of the LFQ intensities revealed by the mass-spectrometry analysis of the IP-Dora^HA^ from OSCs treated (y-axis) and non-treated (x-axis) with bortezomib.

**Figure S3. The cells with *Cul3-KO* and *EloB-KO* are heterozygous on mutation. A.** The expression of *Cul3* and *EloB* in hygromycin-resistance OSC cells upon knock-out. **B, C.** The result of PCR on genomic DNA on the insertion of hyg-containing cassettes into *Cul3* and *EloB* genes. The used primers are numbered and shown as arrows above the *Cul3* and *EloB* genes and *hyg*-cassette sequences (left panel). The result of PCR with indicated primers is shown on the right panel.

**Figure S4. The inducible endogenous knocking down of *Cul3*. A.** The general scheme of the Cu^2+^-induced knockdown of *Cul3*. The constructed cassette MT-shCul3-bsd containing Cu^2+^- activated MT promoter, shCul3, and blasticidin resistance gene *bsd* was integrated into the puromycin resistance gene *PuroR* of genomic pAc-sgRNA-Cas9^Flag^ using Cas9 in OSC^Cas9+^ cells. **B.** The expression level of *Cul3* in the OSCs carrying the genome integrated MT-shCul3-bsd after 72 h of addition of 0 or 1 mM of Cu^2+^ in comparison to the parental OSC^Cas9+^ cell culture. The expression level was measured with qRT-PCR relative to the expression of *rp49*.

**Figure S5. The results of 3’-end mRNA-seq of transcriptomes of *Dora^HA^*and *Dora^HA^-KO* OSCs. A**. The annotation of mRNA-seq reads (UMI counts) on protein-coding and non-protein-coding genes. **B.** The frequency of mapping of reads (cpm of UMI counts) along the metagene. Almost all reads are derived from 3’UTR. **C.** The example of mapping the reads (cpm of UMI counts) on the transcripts of *Ago1*. The peaks of reads correspond to the alternative poly-A sites.

**Figure S6. The Western-blot on Cut in *Dora^HA^* and *Dora^HA^-KO* cells.**

**Table S1.** The primers for the cloning and PCR, probes for the Northern-blot analysis, and adapters for mRNA-seq and miRNA-seq.

**Table S2.** The identified *cis*-NAT-siRNA loci in OSCs.

**Table S3.** The results of the mass-spectrometry of the immunoprecipitates of the cellular lysates of *Dora^HA^* OCSs with anti-HA.

**Table S4.** The differentially expressed genes revealed with mRNA-seq in *Dora^HA^-KO* relative to *Dora^HA^* OSCs.

## Notes

### Competing Interest Statement

The authors have declared no competing interest.

